# Making the *brain-activity-to-information* leap using a novel framework: Stimulus Information Representation (SIR)

**DOI:** 10.1101/658682

**Authors:** Philippe G. Schyns, Robin A.A. Ince

**Affiliations:** Institute of Neuroscience and Psychology University of Glasgow; Institute of Neuroscience and Psychology, University of Glasgow, Scotland G12 8QB, United Kingdom; School of Psychology, University of Glasgow, Scotland G12 8QB, United Kingdom

## Abstract

A fundamental challenge in neuroscience is to understand how the brain processes information. Neuroscientists have approached this question partly by measuring brain activity in space, time and at different levels of granularity. However, our aim is not to discover brain activity *per se*, but to understand the processing of information that this activity reflects. To make this *brain-activity-to-information* leap, we believe that we should reconsider brain imaging from the methodological foundations of psychology. With this goal in mind, we have developed a new data-driven framework, called Stimulus Information Representation (SIR), that enables us to better understand how the brain processes information from measures of brain activity and behavioral responses. In this article, we explain this approach, its strengths and limitations, and how it can be applied to understand how the brain processes information to perform behavior in a task.

“It is no good poking around in the brain without some idea of what one is looking for. That would be like trying to find a needle in a haystack without having any idea what needles look like. The theorist is the [person] who might reasonably be asked for [their] opinion about the appearance of needles.” HC Longuet-Higgins, 1969.

---

Much of human cognition starts with a brain that categorizes stimulus information to behave adaptively^1–3^. Thus, human brains are compulsive categorizers that use stimulus information to perform different categorization tasks. Consider the street scene shown on the left-hand side of Figure 1. The brain can perform numerous categorization tasks on this image: it can identify the country, city, and street; the houses and shops; the moving and stationary cars, their makes and models, age and condition; the people shopping or those just passing by, as well as their gait, identity, emotion and social interactions; it can also infer the weather, time of day, season, and so on.

**Figure 1.**
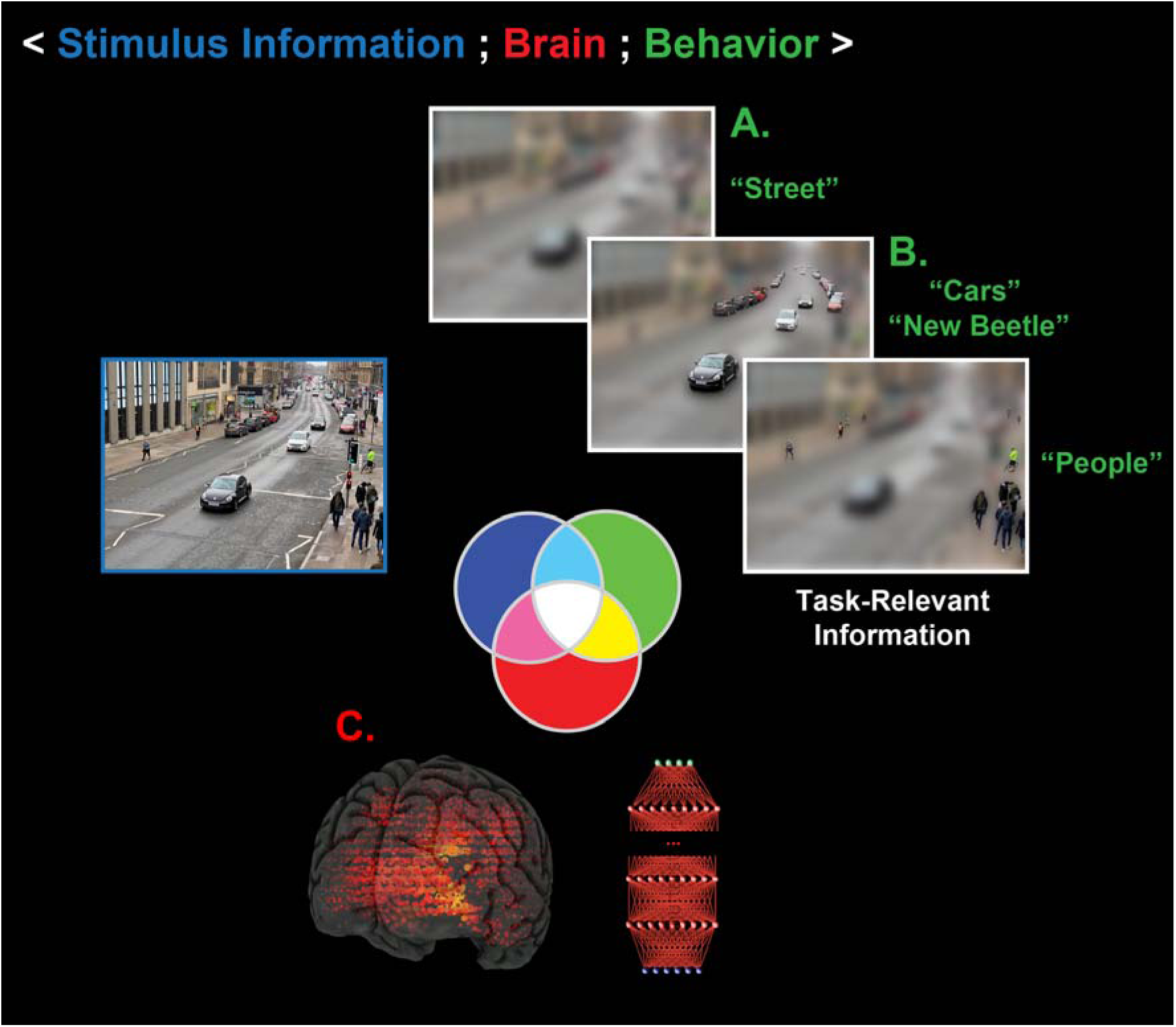
The human brain performs multiple categorization tasks from a single image using task-relevant information. (A) The brain uses coarse, global scene information to categorize this image as “street” in a categorization task. (B) By contrast, local details (i.e. other task-relevant information), as revealed by finer image resolution, support other categorization tasks, such as “cars and their make” and “people”. In the SIR framework, samples of stimulus information (usually in the form of images) are randomly generated and shown to participants to categorize. This approach generates variations in categorization behavior (as represented by the green set). (C) Participants’ brain activity is recorded by neuroimaging techniques (such as EEG and MEG, see the red set) while they perform the task. The three-way interaction between these three SIR components (<stimulus information; brain; behavior>), as represented by the color-coded set of intersections, enables us to better understand how information is processed in the brain.

So, when we record brain activity, we need to circumscribe an experimental task in order to know which task the brain has performed when we analyze its activity, even when a task involves just a single image. And when we circumscribe a task, we still need to characterize the stimulus information that supports a particular categorization. Otherwise, we will not know what information the brain has processed when it categorized the street, or the make of a car, or the identity of a face or its expression, from the same image.

Such task-relevant information processing is a generic, but often neglected theoretical point, that applies both to the interpretation of any sensory categorization in the brain and to its models. For example, Convolutional Neural Networks^4,5^ (CNNs) are brain-inspired, hierarchically organized network models that also use stimulus information to perform multiple categorization tasks, with apparent human-like capabilities. But to realize the promise of CNNs as models of brain information processing^6–12^, a neglected pre-condition needs to be met – that these models process the same task-relevant information as the brain. Otherwise, we cannot explicitly compare how the information processing performed by a CNN with that performed by a human brain.

## The SIR Framework

Stimulus Information Representation (SIR) is a new data-driven framework that can be used to address such issues of information processing by the human brain and by artificial networks. This is because SIR can isolate the specific stimulus information that is processed by brain activity for behavioral responses in a circumscribed task. To do so, SIR uniquely considers the three-way interactions between concurrent trial-by-trial variations of the three main components of an experimental design in the sciences of cognition: stimulus information, behavioral responses and brain activity.

The three components of SIR are gathered and explored in experimental trials. These trials begin by generating random stimulus information to present to participants.

Stimulus information can take different forms. It can consist of images generated by the random selection of pixels^13–16^, or by sampling from generative models of complex stimuli^17–21^. Such randomly generated images (as described in more detail below) are then presented to participants, who are asked to categorize them. So, in this first step, we randomly sample the variables that control stimulus information on each trial (as represented by the blue set on Figure 1). The second component of SIR consists of measuring typical behavioral variables in the performance of a categorization task (including response accuracy, reaction time or confidence ratings) in individual experimental trials (as represented by the green set in Figure 1). When participants perform a categorization task, their behavioral responses to samples of random stimulus information effectively disentangle the stimulus variables that are relevant to that categorization task from those that are not. In this way, the participants’ behavioral responses can reveal which information the brain selectively uses to categorize information as ‘street’, or ‘make of car,’ or ‘face’ and ‘expression’ (see Figure 1). To understand where, when, and how the brain processes these task-relevant variables, we measure its activity while participants perform the categorization task, using various brain-imaging techniques, such as electroencephalography (EEG), magnetoencephalogram (MEG), functional magnetic resonance imaging (fMRI), near infrared spectroscopy (NIRS), electrocorticography (ECoG) or single-cell recordings. The red-shaded information shown in Figure 1 represents variables in brain activity recorded during a task, as recorded by different sensors, at different sources or time points, or representing different neuron firing rates.

Together, the three components of SIR test how randomly sampled stimulus information causally influences brain activity and behavior in a categorization task. In addition, the three-way interactions that occur among these components, as denoted by <stimulus information; brain; behavior>, are represented as the four-color-coded set of intersections shown in Figure 1 (that is, the blue, green, and red sets of the Venn-like diagram, and the white, light blue, magenta, and yellow areas where they intersect). These three-way interactions are unique to SIR. With them, we can address the *brain-activity-to-information* gap with unmatched interpretative precision, as we illustrate in the following sections.

## Applying SIR to understand visual information processing

SIR has been recently applied to analyze brain imaging data collected during the performance of a visual categorization task^22^. This study illustrates how the SIR framework can be used to disentangle brain activity into the processing of task-relevant stimulus information, the processing of task-irrelevant stimulus information, and other brain processes, to enrich the interpretation of brain imaging data. The task that was used is shown in Figure 2A-C. In this task^23,24^, participants were shown an ambiguous image (‘Stimulus’) that can be perceived as being either “the nuns,” or “Voltaire” (squint to see the latter, see Figure 2A). In this task, the green dataset consists of the participants’ behavioral response variable across trials, which can take three possible values: “the nuns,” or “Voltaire,” or “don’t know.” To understand the task-relevant information for each of these responses, we systematically and randomly sampled the image to reveal different pixels to a participant on each trial. The blue dataset shown in Figure 2B, thus includes each image pixel as a distinct stimulus variable (with “on” or “off” values), based on random sampling across trials.

**Figure 2.**
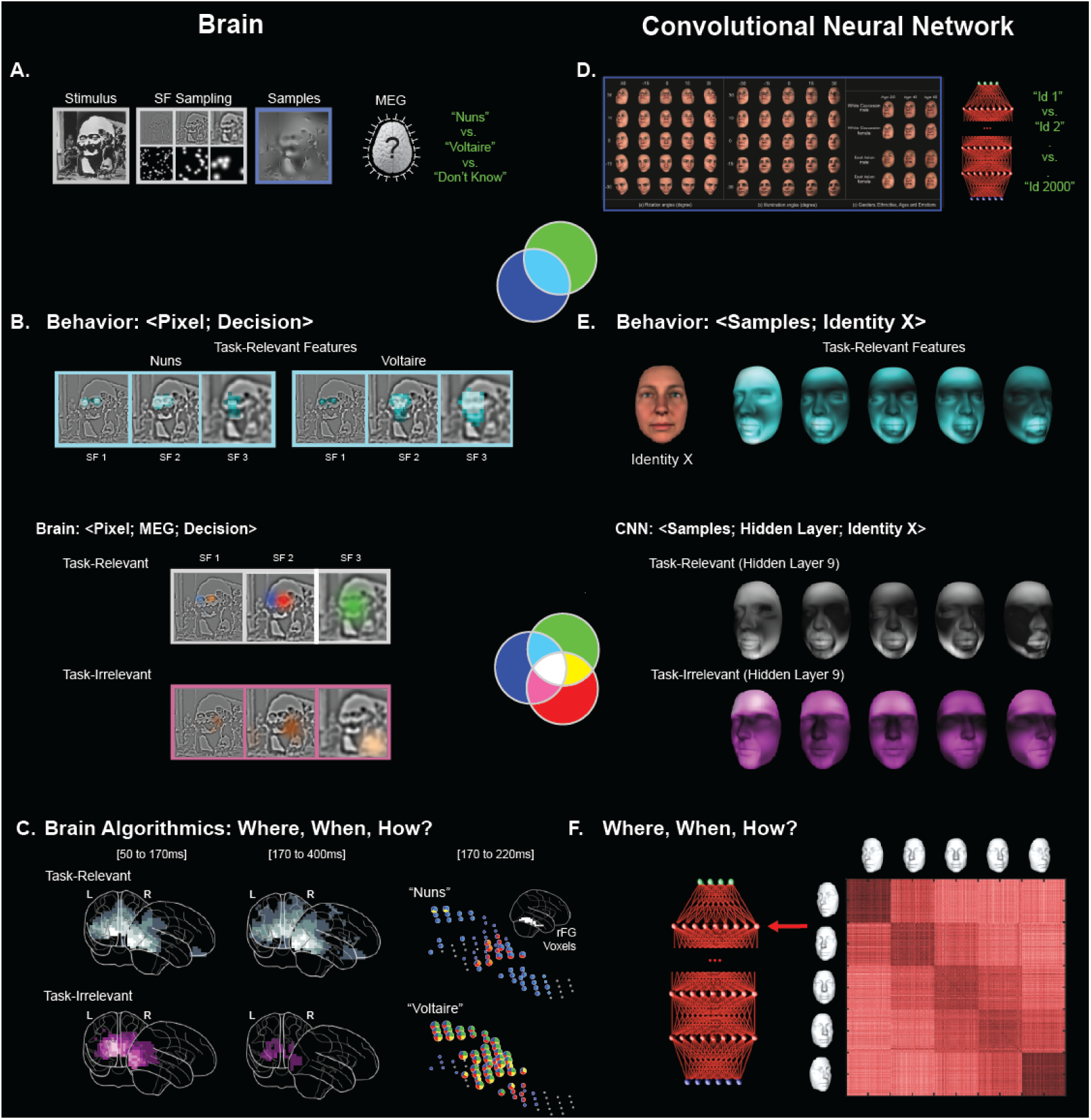
Using SIR to study information processing in the brain and in neural networks. Brain (left column). (A) Pixels of an ambiguous stimulus were randomly sampled across Spatial Frequency (SF) bands using the *Bubbles* procedure^14^. Participants viewed the resulting images and categorize each as being ‘Nuns’, ‘Voltaire’ or ‘don’t know’, while their brain activity (via MEG) and decision behavior were recorded. (B) The pairwise relationship <Pixel; Decision> (see Box 1) for each image pixel was computed to reveal task-relevant (light blue-shaded) features. Three-way relationships (see Box 1) were also computed to reveal task-relevant feature representation (the white triple intersection) and task-irrelevant feature representation (shaded magenta) in brain activity. (C) SIR can be used to show where task-relevant features are represented in the brain, here in the right Fusiform gyrus (rFG) where they are combined into distinct, decision-specific representations. Pie charts standing in for rFG voxels indicate the representational strength of the color-coded task-relevant features framed in white in panel (B). **Convolutional Neural Network (right column)**. (D) 26M facial images were generated using a generative model of face information (see main text for details). 16M images were used to train a 10-layer ResNet and 10M to test its ability to identify faces. (E) Face identification performance saturated at 99.9% correct. The relationship <Sample; Identity X > was computed (see Box 1, and blue intersection) to model the task-relevant features of ‘Identity X’. Across 5 viewpoints, ResNet tracked the same 3D jaw line and mouth shape features. The three-way relationships were also computed to reconstruct task-relevant representations that support ResNet’s identification behavior on layer Y. The white (task-relevant) representations of face shape on Hidden Layer 9 reveal the representation of features for categorization behavior (compare to the blue task-relevant features of ‘Identity X’). Magenta shading shows the task-irrelevant feature representations on Layer 9. (F) Further analyses indicated that the activity of Layer 9 represents the same task-relevant face shape features in a viewpoint-dependent manner.

Using the blue set of randomly sampled image pixels and the green set of corresponding behavioral responses, we then infer the task-relevant stimulus features for each perception, by computing <pixel; behavioral decision>, using (mass-bivariate) pairwise relationships, as represented by the light blue intersection shown on Figure 2 (see Box 1, pairwise relationships). This computation disentangles all image pixels into those that are relevant for task behavior (i.e. for participants to categorize an image as being “the nuns” or “Voltaire”, as represented by the light-blue intersection shown in Figure 2) from those that are not (which are encompassed in the remainder of the blue set).

These light-blue, task-relevant pixels are pivotal for understanding information processing in the brain because they represent the stimulus features that the brain must process to accomplish the behavioral task in question. Task-relevant features are the needles of information that we should search for in the haystack of brain activity. To find these features, we intersect the third component of SIR – in this case, MEG signals, which were recorded during the task. We then compute the overlapping co-representation of the stimulus features into the behavior and brain measures (as <pixel; MEG; decision>, see Box 1 for details). The outcome of this computation identifies the light blue task-relevant features that the red set of MEG variables represent, as highlighted by the white area in Figure 2B, where the three components intersect.

Zhan et al.^22^ used these three SIR components to trace the dynamic flow of task-relevant features that were processed between 50 and 220 ms post stimulus, from their early representation in the visual cortex, through the ventral pathway. In the ventral pathway, we found that task-relevant features converge onto a few MEG voxels at the top of the right fusiform gyrus, ∼200 ms post-stimulus, where they agglomerate into distinct representations that support each behavioral decision (see “task-relevant” images in Figure 2C). Thus, using the three-way interaction between the components of SIR, we traced the dynamics of task-relevant feature processing over the first 220 ms post stimulus, from their early representation in visual cortex to their integration for each behavioral decision in the ventral pathway.

## What do we learn from SIR intersections?

To re-cap, we have three concurrent datasets in the SIR framework – the blue set of stimulus information samples, the red set of brain measures, and the green set of behavioral responses in a task, and their four intersections (colored white, light blue, magenta, and yellow, as shown on Figure 1). The white triple-set intersection is transformative for neuroscience and neuroimaging because it bridges the *brain-activity-to-information* gap by disentangling the different relationships between stimulus, brain activity, and behavior. Specifically, the white set divides each colored intersection into the white component and a remainder. Each of these four intersections contributes its own unique component of interpretation to provide a more detailed understanding of information processing in the brain. We review each intersection in turn.

### The light blue intersection

Within a given task, the light blue remainder set isolates and represents the task-relevant features that the recorded brain measures do not represent. This remainder flags a *de facto* incomplete explanation of the processing of the stimulus information that supports a particular behavior. This is because all task-relevant features should be represented somewhere in the brain to influence behavior. A complete brain measure (such as that captured by a brain-imaging modality) should entirely absorb the light blue remainder (of task-relevant features) into the white set intersection (the light blue remainder is empty in the example shown in Figure 2B-C, where the white framed task-relevant features are all processed in the white brains).

### The magenta and yellow intersections

A magenta remainder reveals task-irrelevant stimulus features, which the brain represents but which do not directly influence behavior in the task (see *Three-way relationships* in Box 1). In Figure 2C, the information processes reduce (i.e. filter out) a travelling wavefront of task-irrelevant feature representations within occipital cortex, around 170 ms post stimulus (see the magenta “task-irrelevant” features and brain in Figure 2B-C). Importantly, although these task-irrelevant features do not influence the participants’ categorization responses, they were amongst the features that were most strongly represented in brain activity in early visual cortex. However, they do not reach the fusiform gyrus in the ventral pathway, as the task-relevant features do.

Finally, the yellow remainder isolates brain activity that relates to a behavior but not to stimulus representation. These brain processes likely reflect other aspects of the task, such as modulation of arousal, response planning, response bias, execution and so forth.

### Using SIR intersections to interpret a CNN

The SIR framework uses each colored set intersection to disentangle a specific kind of relationship between stimulus information, brain activity, and behavior to achieve a finer information processing interpretation of brain activity. SIR can also be used to interpret the processing layers of hierarchically organized CNNs, as shown in Figure 2 (right panel). In a recent study, we taught a 10-layer ResNet architecture to identify 2,000 faces, using 17M images produced by a generative model that varied multiple factors of face appearance. This model employed 25 poses, 25 lighting conditions, random scales and image translations, as well as categorical factors, including three ages, two sexes, two ethnicities, and six facial expressions of emotion plus a neutral expression^25^. We interpreted information processing in ResNet by using SIR and in the same way as we did the brain; by sampling the variables of a generative model of face shape (in the blue set); by recording the variable responses to these image variations by different ResNet layers (in the red set); and by recording the response variations of the real-valued output unit assigned to each identity (in the green set). From these three sets, we then computed the three-way intersection. Figure 2E shows the resulting light-blue task-relevant shape features for Identity X. The white faces reveal the task-relevant features (and their viewpoint-dependent representations) on ResNet Layer 9 (underneath the output layer) that were used to identify Identity X. Magenta faces represent task-irrelevant features on the same Layer 9. Using the SIR framework in this way, we were able to identify task-relevant information and to disentangle its representation in the hidden layers of this CNN from other features irrelevant to the task.

We now turn to some important considerations when applying SIR. As we discuss, the granularity of the sampled information (in the blue set), of the brain measure (in the red set), and of the behavioral response (in the green set) critically determine the rich data-driven interactions that we compute in the four set of intersections, to disentangle information-processing interpretations.

## The Critical Blue Set: What stimulus information should we sample?

As illustrated, the SIR framework depends on a sampling model that creates variations of stimulus information, which in turn cause task-related variations of brain and behavioral responses. For example, in Figure 2A, we sampled pixels across the spatial frequencies of a 2D image. However, it should be clear that the sampling model inevitably constrains the type of stimulus representations that the data-driven framework can discover; for example, the sampling of pixels with Bubbles constrains representations to combinations of contiguous image pixels. This is important because each variable in a sampling model is a *de facto* hypothesis of the stimulus information that the brain represents. Therefore, we are effectively performing mass-trivariate hypothesis testing about the representation of stimulus information variables in brain and behavioral responses.

Consequently, these representational hypotheses determine the structure of the sampled variables to ensure that the stimulus variations tap into the relevant information representations and processes. For example, to study the visual information that supports the prediction of a face from memory, we could set up a model that randomly samples individual pixels to produce a white noise image on each trial (i.e. a sampling model with weak structure, as in^15^). We then instruct the participant that half of the stimuli comprise a face hidden in the noise (when, in fact, there is never such a face) and that their task is to detect it. Computation of the light-blue, task-relevant features characterizes the memory information that serves face prediction, under the constraints of this sampling model^26^. We can address a similar question, using a similar methodology, with the more-sophisticated assumption that memory representations comprise multivariate surface and texture components^27^. In this approach, we would use a structured generative model of multivariate face identity noise (as in the CNN example of Figure 2D-F and ^25,28^) for participants to respond to. We can then study stimulus representations along the occipito-ventral pathway, from the initial projection of the visual input into the laminar layers of early visual cortex to the later task-dependent filtering and representation in the ventral pathway. To do so, we could sample Gabor filters across resolutions (another weakly structured model). Alternatively, we could set up a 3D generative model of these same images to test hypotheses about the faithfulness, compositionality, scale, translation, and rotation invariance of their representations in the ventral pathway. And we could sample both models (i.e. multi-resolution Gabor filters and 3D generative model) simultaneously on each experimental trial.

A sampling model can also be set up to study transformations of representations. For example, while sampling multi-resolution Gabor filters and 3D generative model parameters as described above, we can also include mathematical transformations of the stimulus features (e.g. pooled and normalized Gabor filters; the transfer and integration of surface components) to ascertain the following: which brain regions represent individual variables; which brain regions represent a particular transformation; and how a network of brain regions implements such a transformation. Such sampling of stimulus variables and their transformations can thus support an algorithmic study of the brain^29^, by specifying where, when and how task-relevant variables are represented and transformed in a brain network.

The key point of the critical blue set is that we will not understand the representation and transformation of information that we do not explicitly test. The considerable challenge is to extend stimulus control from low dimensional stimuli (e.g. an oriented Gabor filter at a given spatial frequency) to include generative models that test explicit hypotheses about the task-relevant representations of complex naturalistic stimuli, such as naturalistic faces, objects and scenes, and their transformations across the occipito-ventral pathway^29^.

## Disentangling task-relevant information processing from other processes

In the SIR framework, we randomly sample the variables that control stimulus information on each trial. These variables can affect any measure of behavior (such as accuracy, reaction time, confidence ratings, and so on). They can also affect any measure of brain activity, whether recorded by M/EEG or by 3/7T fMRI, NIRS, ECoG or single-cell recording modalities. And these different imaging modalities record brain activity at different levels of granularity: from individual neurons and population codes^33^; to response characteristics, such as power or phase of oscillatory activity^34^; as well as network interactions^35^, neural synchrony^36^, neuronal firing rates (^37,38^), or spike waves^39^.

In general, to disentangle which variable contributes (or not) to what aspect of information processing, SIR considers the interactions between all combinations of the three classes of variables (see Box 1, *SIR Framework*). Figure 3 illustrates the full potential scope of this effort. At the simplest level (Figure 3A), we have three individual variables (e.g. one individual pixel *X – stimulus*, one response *Y – behavior*, and one voxel *Z* – *brain*), which interact at the four colored set intersections. One level up from this (Figure 3B), we have sets of variables that interact (i.e. *S* image pixels, *N* voxels and *M* behaviors). And finally, we can consider sets of subsets of interacting variables (Figure 3C), where each subset comprises the variables of, for example, a different stimulus sampling model (e.g. Gabor filters or generative models of complex objects), imaging modality or resolution (e.g. EEG or 7TfMRI), or behavior (e.g. reaction time or confidence). At any of these levels, the computations always reduce to three individual variables <*X*_*i*_, *Y*_*j*_, *Z*_*k*_> that interact at the four, colored set intersections. As Box 1 explains, the white triple intersection is quantified with co-information, while the light blue, magenta and yellow remainder double intersections are obtained from conditional mutual information.

**Figure 3.**
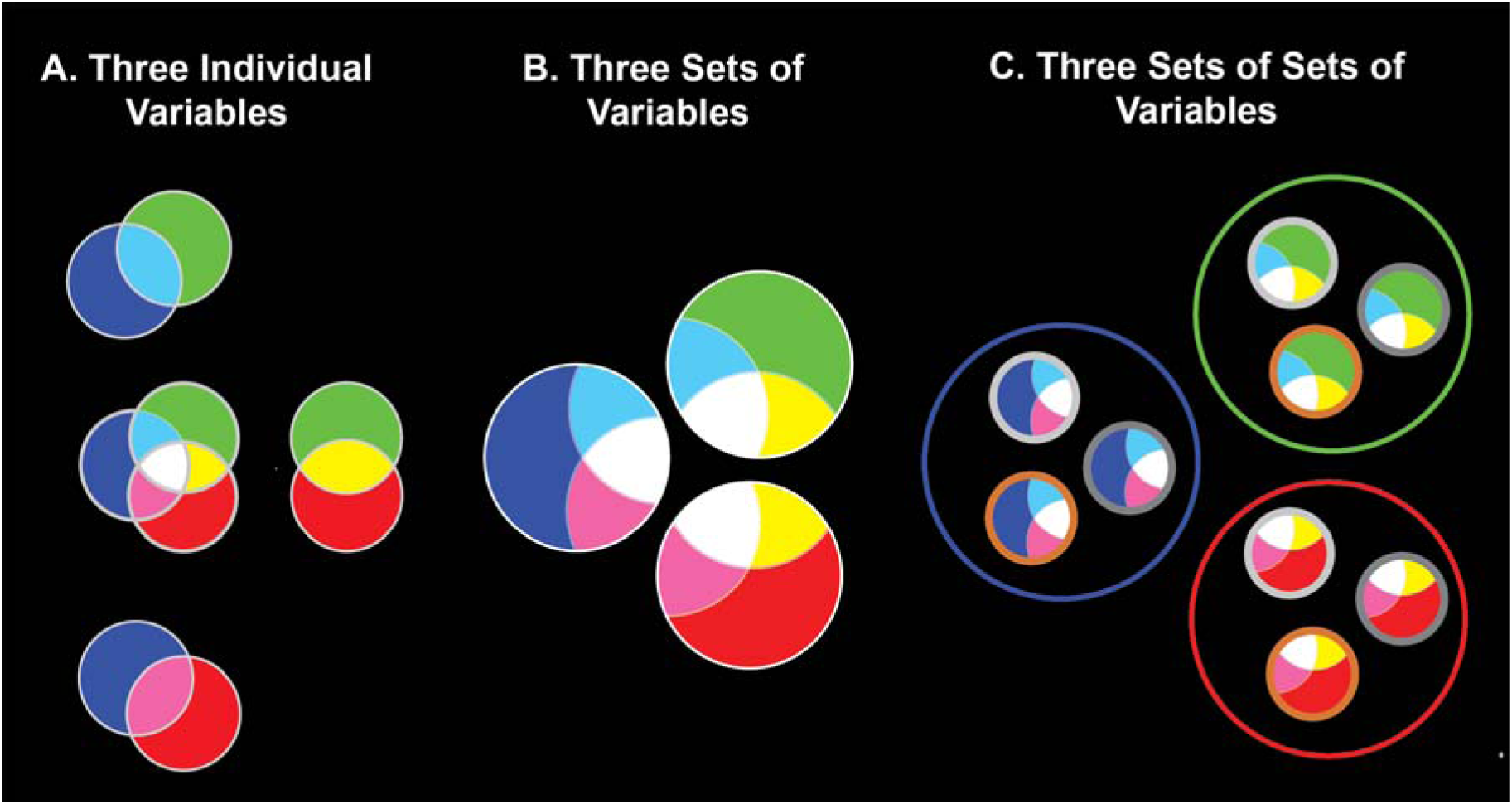
Interactions between SIR variables at different hierarchical levels. *A. Two and Three Individual Variables*. The grey sets represent the entropies of variables *X* and *Y*; their set intersection is mutual information (MI), their shared entropy. With three sets, the blue set represents *H(X)*, the entropy *X* (e.g. one sampled image pixel across trials); the green set *H(Y)*, the entropy of *Y* (e.g. reaction time on an individual trial); the red set represents *H(Z)*, the entropy of *Z* (e.g. one MEG source at a given time point). Co-information (see Box 1) measures the three-way interaction represented as the white set intersection. Conditional MI measures the light-blue, magenta, and yellow remainders of the triple interaction. *B. Three Sets of Variables*. Each set comprises the individual variables of that class considered in the experiment (e.g. *S* stimulus variables; *M* behavioral variables; *N* MEG voxels at multiple time points). These sets are each partitioned for interpretation into the white variables with significant co-information in at least one trivariate combination, the light blue, magenta or yellow variables that have significant conditional MI in at least one trivariate combinations, and the same-color variables that do not interact with any other. *C. Three Sets of Subsets of Variables*. Generalizing, each set comprises multiple modalities as different subsets of variables (e.g. different stimulus sampling models in the blue set, e.g. Gabor sampling variables (e.g. light grey subset) and generative model variables (orange subset); or different modalities or granularities of brain imaging in the red set, such as EEG sensor at different time points (dark grey subset) and fMRI voxels (orange subset)). Across each hierarchical level or subset, the computation depicted in Panel A are applied.

This approach provides a precise, high-dimensional description with which to disentangle task-relevant information processing from other brain processes. To illustrate, consider the Three Sets of Variables in Figure 3B (which structured the examples of Figure 2). Irrespective of the stimulus sampling model, the behavioral response, or the brain measure, the four color-coded interactions of SIR can in principle partition the blue set of stimulus variables into one of four interpretation categories. That is, the stimulus variables that the recorded brain measurements:

1. represent for behavior in the task (these variables are depicted in white in Figure 3B)
2. do not represent but are task-relevant (depicted in light blue)
3. represent but are task-irrelevant (depicted in magenta)
4. do not represent at all (depicted in blue)

Likewise, we can partition the components of brain activity in the red set into the component which:

1. represents the stimulus for behavior (in white)
2. represents the stimulus but not the behavior (in magenta)
3. is involved in other aspects of the task (in yellow)
4. is not involved in the task at all (in the remainder of the red set).

The green set is similarly partitioned into four components. Together, these four intersections provide sufficient precision with which to interpret the data such that we can disentangle the component of brain activity that specifically represents task-relevant information for a given behavior from all other brain activity.

And as mentioned earlier, this framework also extends to multiple modalities, as illustrated in Figure 3C. These richer interactions can provide significant interpretative gains. For example, we could use the white triple intersection to track the representation and transformation of Gabor-generated and generative model-generated task-relevant features in both time and space. Response accuracy, reaction time, and confidence ratings could better dissect those stimulus features that underlie a particular response modality and identify which relationship is better represented in EEG or fMRI modalities. Such an approach could assist the development of richer multi-modal algorithmic network models of task-relevant information processing.

As the combination of interactions explode in each colored intersection, it is worth remembering that the measures of these interactions (i.e. co-Information and conditional MI, see Box 1) provide common scales for comparing, sorting or ranking them, and across any combination of modalities. This implies that we have quantitative (not just qualitative) descriptions to guide the interpretation of the interactions on a common effect-size scale (bits).

## How does SIR compare to alternative frameworks?

The SIR framework is unique because it directly tests and analyses the three-way relationship <stimulus information; brain; behavior>. By contrast, despite recent developments, most statistical techniques in neuroimaging provide bivariate measures of statistical relationships, including highly sophisticated Multivariate Pattern Analysis^40^ (MVPA) and machine learning approaches^41^. For example, encoding and decoding models^42–45^ can quantify a relationship between naturalistic stimuli and multivariate neuroimaging responses, but this is still a measure of bivariate dependence.

Representational Similarity Analysis^46^ (RSA^47^) aims to compare geometries of representations between neural responses and the expected responses of models. However, RSA typically computes an overall average pairwise dissimilarity (e.g. from the mean responses to multiple presentations of a category exemplar) which is not sensitive to the trial-by-trial relationships between the stimulus exemplar and the brain and behavioral responses. Therefore, RSA does not exploit the trial-by-trial variations of the responses and the concurrent interactions between stimulus information, brain measure and behavioral response in a task. When behavior is considered, it is often a perceptual similarity judgment, determined offline, in a separate behavioral experiment, and sometimes even by a different set of participants^48^. However, a recent argument^49,50^ re-emphasized the importance of explicitly including behavior to tease apart the component processes of the brain in neuroscientific explanations. To our knowledge, there is no alternative to the SIR framework to be able to combine the three concurrently recorded variable types to compute their tri-variate interactions during the performance of a specific task.

The SIR framework also highlights ways in which the methodology of brain imaging could be improved. For example, most brain imaging studies of visual categorization use multiple images of various categories (e.g. faces, objects and scenes, or inanimate vs. animate objects) in a one-back or a passive viewing task. However, the brain might not passively perform the explicit categorizations that structure the stimulus categories of our brain imaging experiment. To develop such designs for information processing, we need to add both an explicit control of task behavior (to know the task and the brain’s performance) and an explicit sampling model of stimulus information (to know the task-specific features that the brain must process). Without these, the measured brain activity cannot be unequivocally interpreted as the processing of task-relevant features for that behavior.

These considerations apply equally to the interpretation of CNNs, or to their application as models of the hierarchically organized occipito-ventral pathway. As mentioned earlier, we consider it essential to ensure that the brain and a CNN model are performing the same task in order to compare the brain’s processing of task-relevant information in a categorization task with that of a CNN. The challenge here is to determine that both use the same task-relevant information. Otherwise, you could end up comparing the processing of different visual information across their respective layers. And these different information might appear similar because similarity measures are notoriously under-constrained^51,52^ (until we specify the stimulus features that make two brain measures similar) and context dependent^51,52^ (because the similarity comparison typically involves very few contrast categories). Thus, finding that pairwise representational geometries are similar for brain and CNN responses does not imply that the brain and CNNs are necessarily processing the same information or performing the task in an algorithmically equivalent, or even similar way.

## Conclusion

Using a range of modern imaging modalities, we can now measure the brain activity of an individual participant while they are actively performing an explicit task. During such tasks, stimulus information can be varied on each trial to cause concurrent variations in the participant’s brain activity and behavior. To take full advantage of these richer datasets, we therefore need a new framework that accommodates and exploits these three components – stimulus variation, behavior and brain activity variation. The novel SIR framework does so by considering and computing the interactions between these three sources of trial-by-trial variation. These computations can then be used to disentangle the stimulus, brain activity and response spaces, including at different granularities, depending on the specific experimental design. From these, we make inferences about what information is being processed in the brain for a particular behavior, where and when. SIR can also be applied to study any parametrizable sensory stimulus spaces (e.g. auditory^20,21,53,54^, as well as other cognitive, social and affective tasks (for reviews see^19,55,55,56^, and to study the information processing mechanisms of both brain and in silicon architectures.

We propose, therefore, that the time is ripe to exploit the full capabilities of modern brain imaging technologies and to embrace richer designs that exploit the trial-by-trial trivariate <stimulus information; brain; behavior>. The analyses of such richer designs within the SIR framework can reveal novel interactions that further our understanding of how the brain processes information for behavior.

### BOX 1. The Information Theoretic Underpinnings of the SIR Framework

Since its origin as a mathematical theory of communication in noisy channels^57^, information theory has become a unifying framework for statistics, by providing general measures of the properties of, and the relationships between, probability distributions^58,59^. We use Mutual Information (MI) to measure pairwise relationships between stimulus, behavior and brain variables (represented as the intersections between any two sets in Figure 3), and co-Information^60,61^ (co-I) and conditional MI^62^ to measure the three-way relationships that are unique to precision neuroimaging within SIR (represented as the white intersection and its color-coded remainders in the accompanying figures).

## Pairwise Relationships

The foundational quantity of information theory is entropy, a measure of uncertainty of a random variable *X*, denoted H(*X*), which can be thought of as the information theoretic equivalent of variance. Mutual information (MI) measures the statistical dependence between two variables, *X* and Y, as their shared entropy, *I*(*X*; *Y*) = H(*X*) + H(*Y*) - H(*X, Y*): the sum of the entropies of *X* and *Y* considered as independent variables, minus the entropy of their joint distribution. In Figure 3A, grey sets represent the entropies of H(*X*) and H(*Y*), with MI as their intersection. MI effect sizes are measured in bits, where 1 bit quantifies a reduction in the uncertainty about *Y* by a factor of 2, on average, when observing a value of *X*. These effect sizes are comparable across a range of different tests. For example, the statistical dependence between the sampled stimulus and the participants behavior is quantified on the same effect size scale as the dependence between the stimulus and the recorded neuroimaging responses.

## Three-way relationships

To quantify the relationship between three variables, we use co-information (co-I), denoted I(X; *Y*; *Z*) ^60,61^ the set intersection of three entropies. Co-I is symmetric in the three variables and can be expressed as the intersection of any two MI values, e.g. I(X; Z) + I(Y; Z) - I(lX, YJ; Z). Co-I is the information about Z that is common to *X* and *Y* (or equivalently about *X* common to *Y*, Z, or about Y common to X, Z). For example, in Figure 2, the white triple intersection measures *I*(*pixel*; *MEG*; *decision*), the *pixel* MI that is common to *MEG* and *decision* responses. A positive co-I quantifies *redundancy* between the variables, i.e. they overlap (e.g. the co-representation of the pixel in the MEG and behavior). Negative co-I quantifies *synergy*, when the trial-by-trial relationship between *X* and *Y* provides information about *Z* that is not available from *X* and *Y* considered independently. Like MI, positive and negative co-I also provide effect sizes on common scales for comparison. For example, we can test which of *I*(*pixel*; *MEG*; *decision*) and *I*(*pixel*; *fMRI*; *decision*) is greater, to ascertain which measurement modality (or granularity) better represents in the brain the pixel variations that drive the behavioral decision.

Conditional mutual information (CMI) quantifies the remainder intersections, e.g. I(X; Y|Z), the MI between variables *X* and *Y* conditioned on *Z*. These trivariate quantities are unique to SIR. They depend on the full joint distribution of the three experimental variables and cannot be obtained by considering two variables in isolation, as is the norm in neuroimaging.

## SIR Framework

We consider *S* different stimulus variables (e.g. pixels), *N* different neural responses (e.g. MEG voxels at different post-stimulus time points), and *M* different behavioral variables (e.g. categorization response, confidence rating), all recorded concurrently while a participant performs an explicit categorization task (see Figure 3C). In SIR, for each of the *S × N × M* combinations of stimulus, brain and behavioral variables, we calculate 4 trivariate information quantities by evaluating <stimulus information; brain; behavior> (i.e the white co-Information and the light-blue, magenta and yellow conditional MI, see *Three-way relationship* above and Figure 3A). This *mass-trivariate* computation results in 4 different information processing interaction maps across the full combinatorial space of the three subsets of variables. These maps provide a rich high-dimensional description of the full spatio-temporal properties of stimulus information processing in the brain during the behavioral task. To visualize the results in detail, we can consider one of the three variable classes, by summarizing over all variables of the other two, e.g. by taking the maximum value, or registering statistical significance across any of the possible triples. For example, we can classify each pixel according to the type of interactions that it takes part in, across all neural responses and for any behavioral response. In this way, we can visualize maps of key quantities of interest, as Figure 2C shows, by tracing the spatio-temporal evolution of task-relevant feature representations, where the magenta and white brains are such marginalizations over the task-relevant and task-irrelevant features. In general, the scientific question will dictate the task, the variables within each set of the SIR framework (cf. Figure 3), the specific interactions examined in the 4 interaction maps, and the summary descriptive visualizations. In SIR, the richness of the stimulus information processing description arises from the four unique statistical analyses of all combinations of concurrent tri-variate experimental samples <stimulus information; brain; behavior>.

## References

1. Mervis, C. B. & Rosch, E. Categorization of Natural Objects. Annual Review of Psychology 32, 89–115 (1981).

2. Smith, E.E. & Medin, D.L. Categories and Concepts. (Harvard University Press, 1981).

3. Murphy, G. L. The big book of concepts. (MIT Press, 2002).

4. Hinton, G. E. Learning multiple layers of representation. Trends in Cognitive Sciences 11, 428–434 (2007).

5. LeCun, Y., Bengio, Y. & Hinton, G. Deep learning. Nature 521, 436–444 (2015).

6. Khaligh-Razavi, S.-M. & Kriegeskorte, N. Deep Supervised, but Not Unsupervised, Models May Explain IT Cortical Representation. PLOS Computational Biology 10, e1003915 (2014).

7. Yamins, D. L. K. et al. Performance-optimized hierarchical models predict neural responses in higher visual cortex. PNAS 111, 8619–8624 (2014).

8. Güçlü, U. & Gerven, M. A. J. van. Deep Neural Networks Reveal a Gradient in the Complexity of Neural Representations across the Ventral Stream. J. Neurosci. 35, 10005–10014 (2015).

9. Eickenberg, M., Gramfort, A., Varoquaux, G. & Thirion, B. Seeing it all: Convolutional network layers map the function of the human visual system. NeuroImage 152, 184–194 (2017).

10. Horikawa, T. & Kamitani, Y. Generic decoding of seen and imagined objects using hierarchical visual features. Nature Communications 8, 15037 (2017).

11. Cadena, S. A. et al. Deep convolutional models improve predictions of macaque V1 responses to natural images. PLOS Computational Biology 15, e1006897 (2019).

12. Cichy, R. M. & Kaiser, D. Deep Neural Networks as Scientific Models. Trends in Cognitive Sciences 23, 305–317 (2019).

13. Ahumada, A. & Lovell, J. Stimulus Features in Signal Detection. The Journal of the Acoustical Society of America 49, 1751–1756 (1971).

14. Gosselin, F. & Schyns, P. G. Bubbles: a technique to reveal the use of information in recognition tasks. Vision Research 41, 2261–2271 (2001).

15. Gosselin, F. & Schyns, P. G. Superstitious Perceptions Reveal Properties of Internal Representations. Psychol Sci 14, 505–509 (2003).

16. Murray, R. F. Classification images: A review. Journal of Vision 11, 2–2 (2011).

17. Olman, C. & Kersten, D. Classification objects, ideal observers & generative models. Cognitive Science 28, 227–239 (2004).

18. Greene, M. R., Botros, A. P., Beck, D. M. & Fei-Fei, L. Visual Noise from Natural Scene Statistics Reveals Human Scene Category Representations. arXiv:1411.5331 [cs] (2014).

19. Jack, R. E. & Schyns, P. G. Toward a Social Psychophysics of Face Communication. Annual Review of Psychology 68, 269–297 (2017).

20. Arias, P., Belin, P. & Aucouturier, J.-J. Auditory smiles trigger unconscious facial imitation. Current Biology 28, R782–R783 (2018).

21. Ponsot, E., Burred, J. J., Belin, P. & Aucouturier, J.-J. Cracking the social code of speech prosody using reverse correlation. PNAS 115, 3972–3977 (2018).

22. Zhan, J., Ince, R. A. A., van Rijsbergen, N. & Schyns, P. G. Dynamic Construction of Reduced Representations in the Brain for Perceptual Decision Behavior. Current Biology 29, 319–326.e4 (2019).

23. Smith, M. L., Gosselin, F. & Schyns, P. G. Perceptual Moments of Conscious Visual Experience Inferred from Oscillatory Brain Activity. PNAS 103, 5626–5631 (2006).

24. Ince, R. A. A. et al. Tracing the Flow of Perceptual Features in an Algorithmic Brain Network. Scientific Reports 5, 17681 (2015).

25. Xu, T., Garrod, O., Scholte, S. H., Ince, R. & Schyns, P. G. Using Psychophysical Methods to Understand Mechanisms of Face Identification in a Deep Neural Network. CVPR 9 (2018).

26. Smith, M. L., Gosselin, F. & Schyns, P. G. Measuring Internal Representations from Behavioral and Brain Data. Current Biology 22, 191–196 (2012).

27. Zhan, J., Garrod, O. G. B., Rijsbergen, N. van & Schyns, P. Someone Like You? Modelling Face Memory Reveals Task-generalizable Representations. (2018). doi:10.31234/osf.io/wsvu8

28. Xu, T. et al. Deeper Interpretability of Deep Networks. arXiv:1811.07807 [cs] (2018).

29. Schyns, P. G., Gosselin, F. & Smith, M. L. Information processing algorithms in the brain. Trends in Cognitive Sciences 13, 20–26 (2009).

30. Yuille, A. & Kersten, D. Vision as Bayesian inference: analysis by synthesis? Trends in Cognitive Sciences 10, 301–308 (2006).

31. Zhu, S.-C. & Mumford, D. A Stochastic Grammar of Images. CGV 2, 259–362 (2007).

32. Clark, A. Whatever next? Predictive brains, situated agents, and the future of cognitive science. Behavioral and Brain Sciences 36, 181–204 (2013).

33. Pouget, A., Dayan, P. & Zemel, R. Information processing with population codes. Nature Reviews Neuroscience 1, 125–132 (2000).

34. Schyns, P. G., Thut, G. & Gross, J. Cracking the code of oscillatory activity. PLoS Biol 9, e1001064 (2011).

35. Morcom, A. M. & Friston, K. J. Decoding episodic memory in ageing: A Bayesian analysis of activity patterns predicting memory. NeuroImage 59, 1772–1782 (2012).

36. Singer, W. Neuronal synchrony: a versatile code for the definition of relations? Neuron 24, 49–65 (1999).

37. Ringach, D. L., Hawken, M. J. & Shapley, R. Dynamics of orientation tuning in macaque primary visual cortex. Nature 387, 281 (1997).

38. Popivanov, I. D., Schyns, P. G. & Vogels, R. Stimulus features coded by single neurons of a macaque body category selective patch. PNAS 113, E2450–E2459 (2016).

39. VanRullen, R. & Thorpe, S. J. Surfing a spike wave down the ventral stream. Vision research 42, 2593–2615 (2002).

40. Haxby, J. V., Connolly, A. C. & Guntupalli, J. S. Decoding Neural Representational Spaces Using Multivariate Pattern Analysis. Annual Review of Neuroscience 37, 435–456 (2014).

41. Baker C, Gerven M & Davatzikos C. New advances in encoding and decoding of brain signals. Neuroimage 180, 1–334 (2018).

42. Naselaris, T., Kay, K. N., Nishimoto, S. & Gallant, J. L. Encoding and decoding in fMRI. NeuroImage 56, 400–410 (2011).

43. Weichwald, S. et al. Causal interpretation rules for encoding and decoding models in neuroimaging. NeuroImage 110, 48–59 (2015).

44. Holdgraf, C. R. et al. Encoding and Decoding Models in Cognitive Electrophysiology. Front. Syst. Neurosci. 11, (2017).

45. Kriegeskorte, N. & Douglas, P. K. Interpreting Encoding and Decoding Models. arXiv:1812.00278 [q-bio] (2018).

46. Kriegeskorte, N., Mur, M. & Bandettini, P. Representational Similarity Analysis – Connecting the Branches of Systems Neuroscience. Front Syst Neurosci 2, (2008).

47. Cichy, R. M., Kriegeskorte, N., Jozwik, K. M., van den Bosch, J. J. F. & Charest, I. The spatiotemporal neural dynamics underlying perceived similarity for real-world objects. NeuroImage 194, 12–24 (2019).

48. de-Wit, L., Alexander, D., Ekroll, V. & Wagemans, J. Is neuroimaging measuring information in the brain? Psychon Bull Rev 23, 1415–1428 (2016).

49. Krakauer, J. W., Ghazanfar, A. A., Gomez-Marin, A., MacIver, M. A. & Poeppel, D. Neuroscience Needs Behavior: Correcting a Reductionist Bias. Neuron 93, 480–490 (2017).

50. Tversky, A. Features of similarity. Psychological Review 84, 327–352 (1977).

51. Medin, D. L., Goldstone, R. L. & Gentner, D. Respects for similarity. Psychological review 100, 254 (1993).

52. Owen Brimijoin, W., Akeroyd, M. A., Tilbury, E. & Porr, B. The internal representation of vowel spectra investigated using behavioral response-triggered averaging. The Journal of the Acoustical Society of America 133, EL118–EL122 (2013).

53. Burred, J. J., Ponsot, E., Goupil, L., Liuni, M. & Aucouturier, J.-J. CLEESE: An open-source audio-transformation toolbox for data-driven experiments in speech and music cognition. PLOS ONE 14, e0205943 (2019).

54. Todorov, A., Dotsch, R., Wigboldus, D. H. J. & Said, C. P. Data-driven Methods for Modeling Social Perception. Social and Personality Psychology Compass 5, 775–791 (2011).

55. Dotsch, R. & Todorov, A. Reverse Correlating Social Face Perception. Social Psychological and Personality Science 3, 562–571 (2012).

56. Shannon, C. E. A mathematical theory of communication, Bell Syst. The Bell Systems Technical Journal 27, 379–423 (1948).

57. Cover, T. M. & Thomas, J. A. Elements of information theory. (Wiley New York, 1991).

58. Ince, R. A. A. et al. A statistical framework for neuroimaging data analysis based on mutual information estimated via a gaussian copula. Hum. Brain Mapp. 38, 1541–1573 (2017).

59. McGill, W. J. Multivariate information transmission. Psychometrika 19, 97–116 (1954).

60. Bell, A. J. The co-information lattice. 4th International Symposium on Independent Component Analysis and Blind Signal Separation (ICA2003), Nara, Japan 921–926 (2003).

61. Ince, R. A. A., Mazzoni, A., Bartels, A., Logothetis, N. K. & Panzeri, S. A novel test to determine the significance of neural selectivity to single and multiple potentially correlated stimulus features. Journal of Neuroscience Methods 210, 49–65 (2012).

